# Maraviroc inhibits SARS-CoV-2 multiplication and s-protein mediated cell fusion in cell culture

**DOI:** 10.1101/2020.08.12.246389

**Authors:** Kenneth H. Risner, Katie V. Tieu, Yafei Wang, Michael Getz, Allison Bakovic, Nishank Bhalla, Steven D. Nathan, Daniel E. Conway, Paul Macklin, Aarthi Narayanan, Farhang Alem

**Affiliations:** Center for Infectious Disease Research, George Mason University, Manassas, Virginia, United States of America; Department of Biomedical Engineering, Virginia Commonwealth University, Richmond, Virginia, United States of America; Intellegent Systems Engineering, Indiana University, Bloomington, Indiana, United States of America; Advanced Lung Disease and Lung Transplant Program, Inova Fairfax Hospital, Fairfax, Virginia, United States of America; American Type Culture Collection, Manassas, Virginia, United States of America

**Author notes:** Correspondence: Dr. Aarthi Narayanan.

**Keywords:** SARS-CoV-2, Maraviroc, FDA approved small molecules and antibodies, viral multiplication, cell fusion, predictive modeling

## Abstract

In an effort to identify therapeutic intervention strategies for the treatment of COVID-19, we have investigated a selection of FDA-approved small molecules and biologics that are commonly used to treat other human diseases. A investigation into 18 small molecules and 3 biologics was conducted in cell culture and the impact of treatment on viral titer was quantified by plaque assay. The investigation identified 4 FDA-approved small molecules, Maraviroc, FTY720 (Fingolimod), Atorvastatin and Nitazoxanide that were able to inhibit SARS-CoV-2 infection. Confocal microscopy with over expressed S-protein demonstrated that Maraviroc reduced the extent of S-protein mediated cell fusion as observed by fewer multinucleate cells in the context of drug-treatment. Mathematical modeling of drug-dependent viral multiplication dynamics revealed that prolonged drug treatment will exert an exponential decrease in viral load in a multicellular/tissue environment. Taken together, the data demonstrate that Maraviroc, Fingolimod, Atorvastatin and Nitazoxanide inhibit SARS-CoV-2 in cell culture.

## Introduction

Severe Acute Respiratory Syndrome coronavirus 2 (SARS-CoV-2) is a novel coronavirus which is the causative agent of the coronavirus disease 2019 (COVID-19) pandemic (Zhou et al., 2020). As of November 2021, there are 254,036,628 confirmed cases of COVID-19 resulting in 5,111,176 deaths worldside (Johns Hopkins Coronavirus Resource Center, (Dong et al., 2020)). Coronaviruses (CoV) are generally the cause of common colds resulting in mild respiratory infections (Weiss and Leibowitz, 2011).The emergence of pathogenic coronavirus (CoV) strains with high mortality and morbidity has occurred multiple times over the past two decades, (De Wit et al., 2016). Severe acute respiratory virus (SARS) emerged in 2002 (De Wit et al., 2016; Stadler et al., 2003; Zhong and Wong, 2004). Several SARS-like coronaviruses have been identified in bat species (Banerjee et al., 2019; Ge et al., 2013; Menachery et al., 2015; Wang et al.,2006). Middle Eastern respiratory virus was detected in 2012 (De Wit et al., 2016; Mackay and Arden, 2015). In December of 2019, SARS-CoV-2 began circulating in Wuhan, China resulting in the current COVID-19 pandemic. SARS-CoV-2 is known to be transmitted from human-to-human with inhalational acquisition being the primary mode of transfer (Liu and Zhang, 2020; Nishiura et al., 2020; Song et al., 2020).

Coronaviruses belong to the superfamily *Nidovirales* and subfamily *Orthocoronavirinae* (Weiss and Leibowitz, 2011). CoVs are enveloped, nonsegmented, positive-sense single-stranded RNA viruses (Weiss and Leibowitz, 2011). They are classified in four sub-groups: alpha, beta, gamma and delta. The seven viruses that are known to infect humans belong to the alpha and beta subgroups. HCoV-229E and HCoV-NL62 are classified as alpha while HCoV-OC43, HCoV-HKU1, MERS-CoV, SARS-CoV, and SARS-CoV-2 are beta CoVs. The virus measures approximately 65-125nm in diameter, and the viral genome measures approximately 29.9 Kb (Astuti and Ysrafil, 2020). The viral genome encompasses 14 open reading frames (ORFs) that encode both structural and nonstructural viral proteins. Among the structural proteins, S-protein has achieved significant attention due to the critical role it plays in the interaction of the virus with the ACE2 receptor on host cells (Ahmed et al., 2020). In addition to ACE2, the type II transmembrane serine protease (TMPRSS2) is also required for SARS-CoV-2 entry into cells, thus making these two membrane-associated proteins as the primary determinants for viral entry. ACE2 receptor expression can be detected in various organs besides in the lungs, including heart, kidneys and the gastrointestinal tract. Notably, COVID-19 is characterized by disease manifestations that impact all of the ACE2 positive organs and tissues. The critical aspects of interactions between the viral spike protein and the host membrane proteins ACE2 and TMPRSS2 have led to several therapeutic candidates that interfere with this virus-host protein interactions.

Infection by coronaviruses such as infectious bronchitis virus (IBV) is known to result in cell-cell fusion and formation of large, multinucleated cells referred to as syncytia (Fehr and Perlman, 2015; Sisk et al., 2018). Newly synthesized S-protein either in the context of coronavirus infected cells or cells that over express S-protein, is said to accumulate on the plasma membrane (Lontok et al., 2004).Such S-protein enriched sections of plasma membranes can fuse resulting in cell-cell fusion. Inhibition of IBV infected cells with Abl kinase inhibitors (Imatinib) resulted in reduced syncytia in addition to decreasing viral load (Sisk et al., 2018).This inhibition of S-protein mediated cell fusion by Imatinib could be achieved even in the absence of other viral proteins suggesting that the cell-cell fusion event in coronavirus infected cells is likely to be dependent only on S-protein function. The cell-cell fusion event mediated by coronavirus S-protein is suggested to be controlled by different host enzymatic components than those that influence virus-host membrane fusion. S-protein dependent cell-cell fusion was shown to be independent of cathepsin L which was essential for virus-cell fusion. A novel leupeptin-sensitive host cell protease activated S-protein dependent cell-cell fusion in target cells expressing high levels of ACE2 in the context of SARS-CoV-1 (Simmons et al., 2011).This mechanism of S-protein mediated cell-cell fusion was implicated in viral spread in the context of SARS-CoV-1 infection and the ability of the virus to evade host humoral immune responses, thus posing an important challenge for antibody-mediated viral control (Glowacka et al., 2011). Increased incidence of S-protein mediated syncytia were observed in the context of SARS-CoV-2 S-protein, as compared to SARS-CoV-1, thus highlighting the need to address S-protein mediated cell-cell fusion in SARS-CoV-2 to control viral spread in the infected host (Xia et al., 2020).

A significant contributor towards the morbidity and mortality associated with COVID-19 is the host inflammatory response, with several pro-inflammatory cytokines known to be involved in the tissue damage sustained due to infection. Cytokine storm is suggested to be a significant cause of organ failure and Acute Respiratory Distress Syndrome (ARDS) (Ye et al., 2020). Several host signaling events including the NFkB pathway, JAK-STAT pathway and IFN pathway cumulatively contribute to the proinflammatory environment in target tissues such as the lungs. Onset of ARDS during the later stages of COVID-19 is associated with poor prognosis and is a direct outcome of inflammatory lung damage (Fanelli et al., 2013). Additional complications that are unique to COVID-19 include microclots in blood vessels that supply blood to the heart, lungs and the brain (Ciceri et al., 2020). This disseminated coagulation contributes to death resulting from strokes and heart complications in addition to ARDS (Bansal, 2020; Trejo-Gabriel-Galán, 2020).

It is becoming increasingly apparent that COVID-19 is a combined outcome of viral multiplication in multiple target sites in the infected individual and various host-based complications resulting from dysregulated thrombotic and inflammatory events. An effective therapeutic regimen will need to address the viral and host-based aspects of the disease in order to be able to decrease the health burden and increase survival of affected individuals. Furthermore, such an approach may be essential to address any potential long-term complications that may arise in survivors, especially those who are considered to be high-risk populations. High-risk individuals include those with preexisting health conditions such as coronary heart disease, and inflammatory states including asthma and diabetes. Seeking a therapeutic regimen that may be easily adapted to such high-risk individuals, we focused our attention on a subset of FDA-approved small molecules and biologics that are commonly used to treat heart disease, inflammation and infectious disease. We screened a total of 22 such molecules and have identified four small molecules (Maraviroc, FTY 720, Atorvastatin and Nitazoxanide) and three antibodies (Bevacizumab, Eculizumab and Anti-CD126/IL6R), all of which were able to decrease infectious virus titers in cell culture. Additional studies performed with the prioritized small molecule inhibitors demonstrated that these compounds moderately impacted viral entry and interfered with the interaction between the spike protein and ACE2 to varying degrees. Cell biology studies demonstrate that Maraviroc was able to decrease the extent of S-protein-mediated cell fusion. Our efforts to develop simulation models of the antiviral outcomes of Maraviroc on a tissue level demonstrated that sustained availability of drug will have a significant impact on decreasing viral load and cellular fusion and increasing cell survival. Simulations also show that drugs targeting viral fusion may increase efficacy of anti-replication drugs when used in combination.

## Material and Methods

### Virus, Cell lines and reagents

SARS-CoV-2 (2019-nCoV/USA-WA1/2020) was obtained from BEI Resources (NR-52281). VERO African green monkey kidney cells (ATCC Cat# CCL-81) were obtained from the American Type Culture Collection. African green monkey kidney cells (Vero) were maintained in DMEM supplemented with 5% heat-inactivated fetal bovine essence (FBE) (VWR Life Sciences 10803-034), 1% Penicillin/Streptomycin (Gibco, 15-140-122) and 1% L-Glutamine (VWR Life Sciences, VWRL0131-0100) at 37 °C and 5% CO2. All reagents for cell maintenance were prewarmed to 37 °C before use.

### Inhibitors and Antibodies

Antibodies Eculizumab (HCA312), Bevacizumab (HCA182) and CD126/IL-6R (AHP2449) were obtained from Biorad.

Dipyridamole (06-915-00), Ciclesonide (63-575), FTY720 (61-761-0), Pirfenidone (P1871100MG), Atorvastatin (A24761G), Indomethacin (1065525G), Bromhexine HCl (B405425G), Maraviroc (37-561-0), Darunavir (67-101-0), Danoprevir (50-190-4862), Azithromycin (37-715-0), Nitazoxanide (N1031200MG) and Tiecoplanin (AC455400010) were obtained from ThermoFisher Scientific.

Valganciclovir (S4050), Rimantadine (S5484), Letermovir (S8873), and Baloxavir (S5952) were obtained from Selleckchem.

### Viral infection and inhibitor treatment

Vero cells were seeded at a density of 3E4 cells/well. The inhibitors were resuspended in dimethyl sulfoxide (DMSO) prior to use and stored as recommended by the manufacturer as stock solutions (5 mM – 100 mM, varied depending dry weight and molecular weight of inhibitor received). Inhibitors were diluted to stated concentrations in cell culture media. Cells were pretreated with the target inhibitors before infection for 2h. Virus was diluted in media to appropriate MOI and placed on cells. The plate was incubated for 1 hour at 37°C, 5% CO2 to allow the uptake of virus. Viral inoculum was removed, cells were gently washed three times with Dulbecco’s phosphate-buffered saline (DPBS) without calcium and magnesium (Gibco 14190144) and conditioned media with the inhibitors was added back onto the cells. Cells were incubated at 37°C and supernatants were collected at indicated times post infection and stored at -80°C until required for further analyses.

### Plaque assay

Plaque assays were performed using Vero cells grown to a cell number of 2E5 cells/well in a 12-well plate. Supernatants from infected cells were serially diluted in media and used to infect Vero cells for one hour. After infection a 1:1 overlay consisting of 2X Eagle’s Minimum Essential Medium without phenol red (Quality Biological, 115-073-101), supplemented with 10% fetal bovine serum (FBS) (Gibco, 10,437,028), non-essential amino acids (Gibco, 11140-050), 1 mM sodium pyruvate (Corning, 25-000-Cl), 2 mM L-glutamine, 20 U/ml of penicillin and 20 μg/ml of streptomycin) and 0.6% agarose was added to each well. Plates were incubated at 37°C for 48 hours. Cells were fixed with 10% formaldehyde for 1 hour at room temperature. Formaldehyde was aspirated and the agarose overlay was removed. Cells were stained with crystal violet (1% CV w/v in a 20% ethanol solution). Viral titer (PFU/mL) of SARS-CoV-2 was determined by counting the number of plaques.

### Cell viability assay

Cell viability was determined using the CellTiterGlo assay (Promega, G7570). Inhibitors were resuspended in DMSO and stored appropriately. Inhibitors were diluted to required concentrations in cell culture media. Vero cells were seeded at a density of 3E4 in white-walled 96-well plates. Inhibitors were overlaid on cells and incubated at 37°C, 5% CO2 for 24 hours. CellTiter-Glo substrate was used according to the manufacturer’s instructions and luminescence was determined using a DTX 880 multimode detector (Beckman Coulter) with an integration time of 100ms/well. Cell viability was determined as percent cell viability normalized to untreated control.

### ACE2: Spike S1-Biotin (SARS-CoV-2) Inhibitor Screening Assay

Small molecules inhibitors were screened for ability to disrupt SARS-CoV-2 spike proteins binding to ACE2 receptors. ACE2: Spike S1-Biotin (SARS-CoV-2) Inhibitor Screening Assay Kit (BPS Bioscience, Cat# 79945) was utilized per manufacturer’s instructions. Plate was washed and blocked in between each addition step with provided wash buffer and blocking buffer. A nickel-coated plate was treated with his-tagged ACE2. Drugs were screened at a low concentration (25 μM) and a high concentration (100 μM). Inhibitor or DMSO was added to the plate to bring final dilution to indicated concentration. Plate was incubated for 1 hour at room temperature with shaking. SARS-CoV-2 spike S1-biotin was added to the plate. Plate was incubated for 1 hour at room temperature with shaking. Streptavidin-HRP is added to the plate. Plate was shaken for one hour at room temperature. HRP substrate was added to the plate and the Chemiluminescence was measured immediately using a DTX 880 multimode detector (Beckman Coulter) with an integration time of 1 second, and delay of 100ms.

### rVSV Pseudotyped SARS-CoV2 Spike Neutralization Assay

The vesicular stomatitis virus (rVSV) with deleted glycoprotein and replacement SARS-CoV-2 spike was acquired from IBT Bioservices. Vero cells were seed at a density of 6E4 in a black, 96-well tissue culture plate. Cells were maintained in EMEM supplemented with 10% fetal bovine serum (FBS) and 1x penicillin/streptomycin (P/S). Cells were incubated overnight 37°C and 5% CO2. Virus was diluted per manufacture’s protocol and incubated with drug at a final concentration of 5 μM or DMSO control in unsupplemented media for one hour at room temperature. Media was removed from plate, replaced with treated or untreated virus. Plate was incubated for one hour at 37°C. EMEM supplemented with 2% FBS and 1x P/S was added to the wells and the plate was incubated overnight at 37°C and 5% CO2. Cells were lysed with 1X Passive Lysis Buffer for 30 minutes while shaking. Firefly Luciferase Assay substrate was added to the plate. Plate was read immediately using a DTX 880 multimode detector (Beckman Coulter) with an integration time of 1 second.

### Transfection Methods

Vero cells were grown on 6-well plates and were transiently transfected with Lipofectamine 2000 (Invitrogen, 11668019) based on the manufacturer’s protocols. In brief, prior to treatment, Lipofectamine 2000/siRNA complexes were prepared in reduced serum medium, OptiMEM (Gibco, 31985088), at the recommended ratio. Cells were treated for 6 hours before being replaced with EMEM complete medium.

### Microscopy analysis of cell-cell fusion

African green monkey kidney (Vero) cell lines (ATCC Cat# CCL-81) were grown in Eagle’s Minimum Essential Medium (EMEM) (Corning) supplemented with 10% heat-inactivated fetal bovine serum (Gibco, USA). Cells were cultured at 37°C in a humidified incubator supplied with 5% CO2. pTwist-EF1alpha-SARS-CoV-2-S-2xStrep (a gift from Nevan Krogan, Addgene plasmid # 141382) was used to express S-protein from the original SARS-CoV-2 strain (WT). To express D614G S-protein this S-protein construct was mutated to introduce the D614G substitution. Additionally, we also expressed WT and D614 S-protein-FLAG (Addgene plasmids 156420 and 156421, gift of Hyeryun Choe) for improved immunostaining. Lipofectamine 2000 (ThermoFisher) was used to transfect cells with the S-protein DNA (per manufacture instructions). Maraviroc (Millipore Sigma, PZ0002) was dissolved in DMSO at a 1 mM (stock). Before or following transfection, 5 μM of Maraviroc solution was added to each well with complete EMEM medium. Vero cells were transferred to 35 mm Glass bottom dishes (Cellvis, D35-20-1-N) 24 hours after transfection. Then cells were fixed with 4% para-formaldehyde (Electron Microscopy Sciences, 15710) in PBS (Gibco, 14040182) 24 hours after attachment and then incubated with PBS. Thereafter, cells were mounted using Prolong Diamond Antifade Kit (Invitrogen, P36970) according to manufacturer’s instructions. Hoechst 33342 (ThermoFisher, H3570) were used to identify nuclei. 2xStrep tagged S-protein was stained using StrepMAB-Classic DY-488 conjugate (IBA Lifesciences, 2-1563-050) and FLAG tagged S-protein was stained using anti-FLAG (clone M2, Sigma-Aldrich, F1804). SARS-CoV-2 Spike Antibody (ProSci, 3525) was also used to detect expressed S-protein. Actin-stain 555 phalloidin (Cytoskeleton, PHDH1-A) was used to label the actin cytoskeleton. Cells were imaged using a Zeiss LSM 710 Laser Scanning Microscope. Images were analyzed by Fiji (U.S. National Institutes of Health, Bethesda, MD). Total nuclei and S-protein transfected nuclei were counted using Cell Counter plugin.

### Simulation modeling

To simulate Maraviroc treatments, we adapted an open source, community-developed simulator (Version 3.2) for SARS-CoV-2 infection dynamics in tissue that currently includes diffusive viral spread through the tissue, ACE2 receptor dynamics, intracellular viral replication, single-cell viral and antiviral responses, and immune response (Getz et al., 2020). The simulator uses an agent-based modeling approach (PhysiCell (Ghaffarizadeh et al., 2018)) to simulate individual epithelial and immune cells. After adapting to the *in vitro* experimental conditions (no immune components), we extended the model to simulate cell fusion under the assumption that infected cells express spike protein on the surface (modeled to be correlate with the infected cell’s assembled virus) that can bind to to nearby cells, leading to cellular fusion (Braga et al., 2021; Stoyanov et al., 2020). While inhibition experiments suggest that fusion is not necessarily driven by binding to ACE2 receptors on nearby cells (see **Results**), we use unoccupied ACE2 receptors in our model to more broadly model possible interaction targets (Hoffmann et al., 2020) to mediate fusion. Thus, we model the probability *P*_fusion_ of two cells fusing as

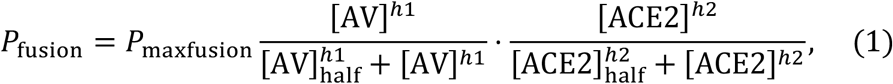

where *h1* and *h2* are Hill powers, [AV] is assembled virus in the infected cell (correlated with surface expression of spike protein), [ACE2] is unoccupied ACE2 receptors on the cell to be fused (proxy model of all possible binding targets), [AV]_half_ and [ACE2]_half_ are respective half-maximum values, and *P*_maxfusion_ is the maximum fusion probability when the cells have plentiful surface expression of spike protein and binding targets. In this model, fused cells continue viral replication with the same mass-action kinetics as individually-fused cells.

To prepare for treatment simualtions, we added a generalized pharmacodynamic response model to each cell agent, where local drug availability could drive a combination of responses (including dose-dependent inhibition of cellular fusion, viral endo- and exocytosis, or viral replication) in each simulated cell. This allows us to model both the specific targeted effects of anti-fusion drugs as well as off-target, secondary, or mixed effects that are typically observed for drugs (e.g., Braga et al. (2021)). See Getz et al. (2020) for full details on the ACE2 receptor binding and trafficking, viral replication and exocytosis, and infected cell death response. See **Figure S1A** for the model overview.

To model the individual-cell pharmacodynamics, we first modeled the rate of drug internalization as diffusion through the cell membrane, allowing us to track the accumulated total drug (*n*) in each cell (Equation 2). Similarly to Aljayyoussi et al. (2017) and Aljayyoussi et al. (2019), we model the total effect *E* for each cell based on the internal drug concentration *ρ*_intra_ using a Hill function (Equation 3). The effect is then used to modulate a behavioral parameter *p* (endocytosis rate, exocytosis rate, replication rate) from its untreated value (*p*_0_) to its value *p*_max_ at maximum drug effect when *E* = 1. (Equation 4). Note that *p*_max =_ *p*_0_ for any parameter not affected by the simulated drug. In addition, we can model the impact of partial drug efficacy by multiplying the effect by a sensitivity *S*, where *S* = 0 denotes a drug with no efficacy (e.g., it fails to bind its target, or binding its target does not affect cell phenotype), and *S* = 1 is an ideal drug that fully binds and inhibits its target. In addition, we used a nonliear interpolation for calculating fusion probability under drug effect (Equation 5). Taken together, we model:

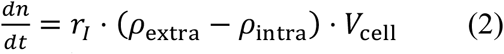

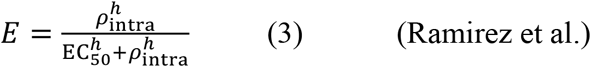

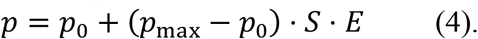

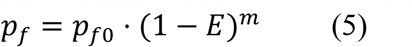

Here, *r*_*I*_ is drug internalization rate, *ρ*_ectra_ and *ρ*_intra_ are the extracellular and intracellular drug concentrations, *V*_cell_ is cell volume, EC_50_ is the drug dose that gives 50% of maximum effect, and *h* is Hill power (Goutelle et al., 2008). *p*_*f0*_ is fusion probability without drug effect (see Equation 1), *m* is the inhibition coefficient.

Full simulation source code is available as open source under the BSD license; see Appendix A. We used *xml2jupyter* (Heiland et al., 2019) to create a cloud-hosted version of this model; interested readers can interactively run the model in a web browser at https://nanohub.org/tools/virion2pbpd). See **Figure S1B-C**.

### Simulation model fitting

We began with the default parameter values from Getz et al. (2020) and adjusted to match viral titer measurements at 16 hours under control conditions and a 5 μM dose of Maraviroc. First, we adjusted the receptor dynamics parameters (ACE2 binding rate, ACE2 endocytosis rate, ACE2 cargo release rate and ACE2 recycling rate), viral replication parameters (virion uncoating rate, RNA preparation rate, viral protein synthesis rate, and virion assembly rate) and export rate, and the fusion probability to match the viral titer in untreated Vero cells at 16 hours, assuming that 3000 cells are initially infected.

To calibrate the drug uptake rate, we looked at prior experiments which show that peak cytoplasmic drug concentrations are reached in about 3.1 hours (300mg dose) (Carter and Keating, 2007). To match this drug uptake time scale, we set the internalization *r*_*I*_ at 0.0116 *μ*M · μ*m*^3^ /min (1 hour to reach half maximum). We initially approximated EC_50_ as 2.5 *μ*M (half of experimental dose), the Hill power *n* as 1, and *m* as 1 as an intial parameter estimate.

We then sought a parsimonious pharmacodynamic model that could fit the full response of the 5 μM Maraviroc dose, with both a reduction in viral titer at 16 hours and cell fusion at 48 hours. First, we attempted to fit the simplest possible model where the drug inhibits cell fusion only but has no effect on viral endocytosis and exocytosis, and no effect on replication. While we were able to fit the model to match the reduction of cell fusion, this model could not match experimental measurements of reduced viral titer. Hypothesizing that a drug that impacts viral protein localization on the cell surface may also impact endocytosis and exocytosis (similarly to other recent observations (Hoffmann et al., 2020)), we next modeled the drug as inhibiting fusion, endocytosis, and exocytosis (with equal impact on endocytosis and exocytosis, for simplicity). To match with both the reduction of viral titer at 16 hours and the inhibtion of cellular fusion at 48 hour with a dose of 5 μM, we varied the Hill power *n* and inhibition coefficient *m* until the full dynamical model yielded a viral titer and cell fusion that matched experimental measurements. The resulting parameters were EC_50_ = 2.5 *μ*M, *n* = 2.5, and *m*=2.5. See Goutelle et al. (2008) for more details on common pharmacodynamic forms.

Because this was the simplest (most parsimonious) set of drug effect assumptions that could fit the data, we used it for the remainder of the study. We note that simulating Maraviroc as partially inhibiting replication as well as endo/exocytosis and fusion could also potentially fit the data, but as a less parsimonious model.

### Computing

We used a total of 13 parameters sets. Because the simulation model is stochastic, we ran 10 simulation replicates for each parameter set. All simulation images are chosen from one representative replicate, and all aggregate dynamical curves (e.g., viral titer versus time) are reported as the mean and confidence interval as 95% over all 10 simulation replicates. The simulations were performed on the Big Red 3 supercomputer at Indiana University. Each simulation was run on a single node using 24 threads. All jobs (130 runs) were submitted as a batch. Each simulation was completed in approximately 25 minutes, and with a total wall time of approximately of 9 hours.

### Statistical Analysis

Statistical analyses were performed using the software Prism 5 (Graph Pad). Data are presented as mean ± standard deviation (SD) after analysis with unpaired, two-tailed t-test or one-way ANOVA. Differences in statistical significance are indicated with asterisks: * *p*<0.05; ** *p*<0.01; *** *p*<0.001; **** *p*<0.0001.

### Data and Code Availability

The simulation source code is available at:

GitHub repository: https://github.com/pc4covid19/pharmacodynamics-submodel

Zenodo archive: https://zenodo.org/badge/latestdoi/282789470

The nanoHUB web version of the model can be accessed at: nanoHUB DOI: http://dx.doi.org/doi.10.21981/CCFV-ES15

## Results

### Identification of anti-SARS-CoV-2 activity of FDA approved small molecules

The investigation was conducted with 19 FDA-approved small molecules that are commonly prescribed to the general population which included antiviral and anti-parasitic compounds. The investigation also included compounds that were documented to alter host signaling cascade events and exhibit anti-inflammatory properties. The list of small molecules that were tested is included in **Figure 1A**. The investigation was conducted using Vero cells and the Washington strain of SARS-CoV-2 (2019-nCoV/USA-WA1/2020). The toxicity of the small molecules was first quantified by treating the cells with the inhibitor at a low, medium and high concentration (1 μM, 5μM and 10 μM) for 24 hours after which cell viability was quantified using Cell Titer Glo assay (**Figure 1B**). Almost all of the small molecules demonstrated less than 10% cytotoxicity in this assay with the exception of FTY720 and Atorvastatin that showed 20% and 10% reduction in cell viability (compared to the untreated control) at the highest concentration. Anti-SARS-CoV-2 activities of these small molecules were quantified by plaque assay and the outcomes are included in **Figure 1C**. Vero cells were pre-incubated with nontoxic concentrations of the compounds for 1 hour prior to infection. Infection with SARS-CoV-2, Isolate USA-WA1/2020 was carried out by incubating the virus with cells (multiplicity of infection [MOI]: 0.1) in the presence of the designated inhibitor for 1 hour. After allowing time for the infection, the virus overlay was removed and media containing the appropriate inhibitor was added back to the cells. DMSO treated cells were maintained as controls alongside. The amount of infectious virus in the supernatants of the drug-treated cells was quantified by plaque assay at 16 hours post infection and compared to the DMSO-treated control.

**Figure 1.**
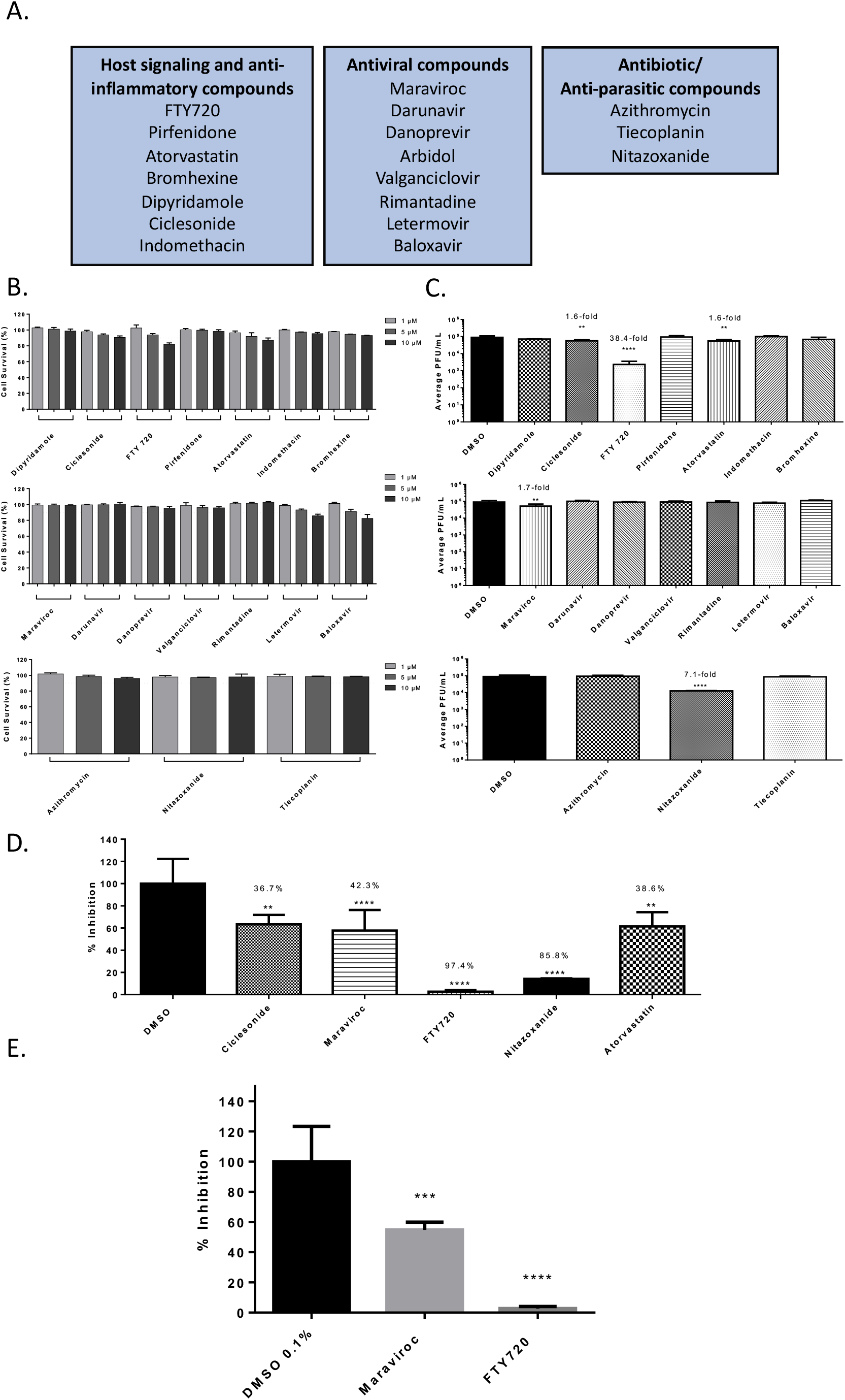
Efficacy of small molecule inhibitors against SARS-CoV-2 in Vero cells 16 hpi with inhibitor in viral inoculum. (A.) Classification of the Small molecule inhibitors that were tested. (B.) Drug toxicity in Vero cells at 24 hours. Cells were seeded at a density of 3E4 and overlaid with media conditioned with 0.1% DMSO or the indicated concentration of inhibitor. Cells were incubated for 24 hours at 37ºC at 5% CO2. Viability was determined using CellTiter Glo assay. (C.) Efficacy of small molecule inhibitors in Vero cells 16 hpi. Cells were seeded at a density of 3E4 and overlaid with conditioned media containing 0.1% DMSO or 5 μM of inhibitor. Cells were incubated for 2 hours. Conditioned media was removed. Viral inoculum (MOI: 0.1) containing 5 μM of inhibitor was added to cells and incubated for 1 hour at 37ºC and 5% CO2. Viral inoculum was removed, cells were washed with DPBS, and conditioned media was returned to the cells. Culture was incubated for 16 hpi. Supernatants were collected and a viral plaque assay was used to quantify the viral titers. (D.) Inhibition of SARS-CoV-2 Washington 2020 observed at 16 hpi, represented as relative percentages. (E.) Inhibition of SARS-CoV-2 Italy 2020 observed at 16 hpi, represented as relative percentages. The quantitative data are depicted as the mean of three biologically independent experiments ± SD. * *p* < 0.05; ** *p* < 0.01; *** *p* < 0.001; \*\*\*\**p* < 0.0001.

Among the antiviral compounds that were tested, Maraviroc demonstrated a 1.7-fold decrease (p<0.0001) in viral load as compared to the DMSO control **(Figure 1C)**. Among the anti-parasitic compounds that were tested, Nitazoxanide demonstrated a 7.1-fold decrease (p<0.0001) when compared to the DMSO control. Interestingly, Nitazoxanide was suggested as a potential anti-SARS-CoV-2 therapeutic based on prior activity observed against MERS-CoV (Martins-Filho et al., 2020). Nitazoxanide was also suggested as a combinatorial therapeutic along with Azithromycin (Kelleni, 2020). Nitazoxanide is currently undergoing clinical trials for safety and efficacy to treat COVID-19 patients (ClinicalTrials.gov Identifier: NCT04435314). In the set of compounds that influenced host signaling and inflammatory events, FTY-720 showed a 38.4-fold decrease (p<0.0001) of SARS-CoV-2 viral titer as compared to the DMSO control. FTY-720 is FDA-approved for the treatment of multiple sclerosis and is documented to exhibit antiinflammatory activities by modulating the NFkB signaling cascade (Pul et al., 2016). In this class of compounds, Atorvastatin, a commonly used statin for the treatment of high cholesterol, was noted to decrease viral load 1.6-fold (p<0.01) as compared to the DMSO control. The relative percent inhibitions of these prioritized compounds is indicated in **Figure 1D**, in comparison with the DMSO control. Maraviroc and FTY-720 were screened against SARS-CoV-2, Isolate Italy-INMI1 **(Figure 1E)** in addition to Isolate USA-WA1/2020.Maraviroc decreased viral titers 45.2% while FTY-720 showed a 97.4% decrease.

### Identification of anti-SARS-CoV-2 activity of FDA approved biologics

Three FDA-approved biologics namely, Bevacizumab, Eculizumab and CD126/IL6R mAb were evaluated for anti-SARS-CoV-2 activity in a similar manner as described above (**Figure 2A**). Bevacizumab (Avastin) is an inhibitor of VEGF-R2 by binding to VEGF-A and is used for the treatment of several types of cancers (Mahfouz et al., 2017; Papachristos et al., 2019). Eculizumab (Soliris) is undergoing clinical trials for use as a therapeutic for the treatment of COVID-19 patients, potentially for the anti-inflammatory activities and the ability to moderate inflammation related complications (ClinicalTrials.gov Identifier: NCT04288713). Briefly, toxicity of the biologics was evaluated by incubating Vero cells with a low (1 μg) and high (2.5 μg) concentration of the target biologic for 24 hours after which cell viability was quantified by Cell Titer Glo (**Figure 2B**). Less than 10% cytotoxicity was detected for these molecules at the concentrations tested, with the exception of CD126/IL-6R antibody. A concentration of 2.5 μg was selected for efficacy evaluation for the less toxic biologics and 1 μg was selected for the CD126/IL-6R antibody. Impact of inhibitor treatment on SARS-CoV-2 infectivity was assessed by performing infection of Vero cells (MOI: 0.1) in the presence of the biologic at nontoxic concentration for 1 hour. The infected cells were incubated with the biologic for an additional 16 hours after which infectious virus load in the supernatant was quantified in comparison with the DMSO control (**Figure 2C**). Bevacizumab and Eculizumab treatment resulted in a 4.2-fold decrease (p<0.0001) and a 3.7-fold decrease of virus titers, respectively, which coorilate to 76.2% and 73.1% total reduction in virus respectively **(Figure 2D)**. CD126/IL6R antibody resulted in 1.9-fold decrease of virus load.

**Figure 2.**
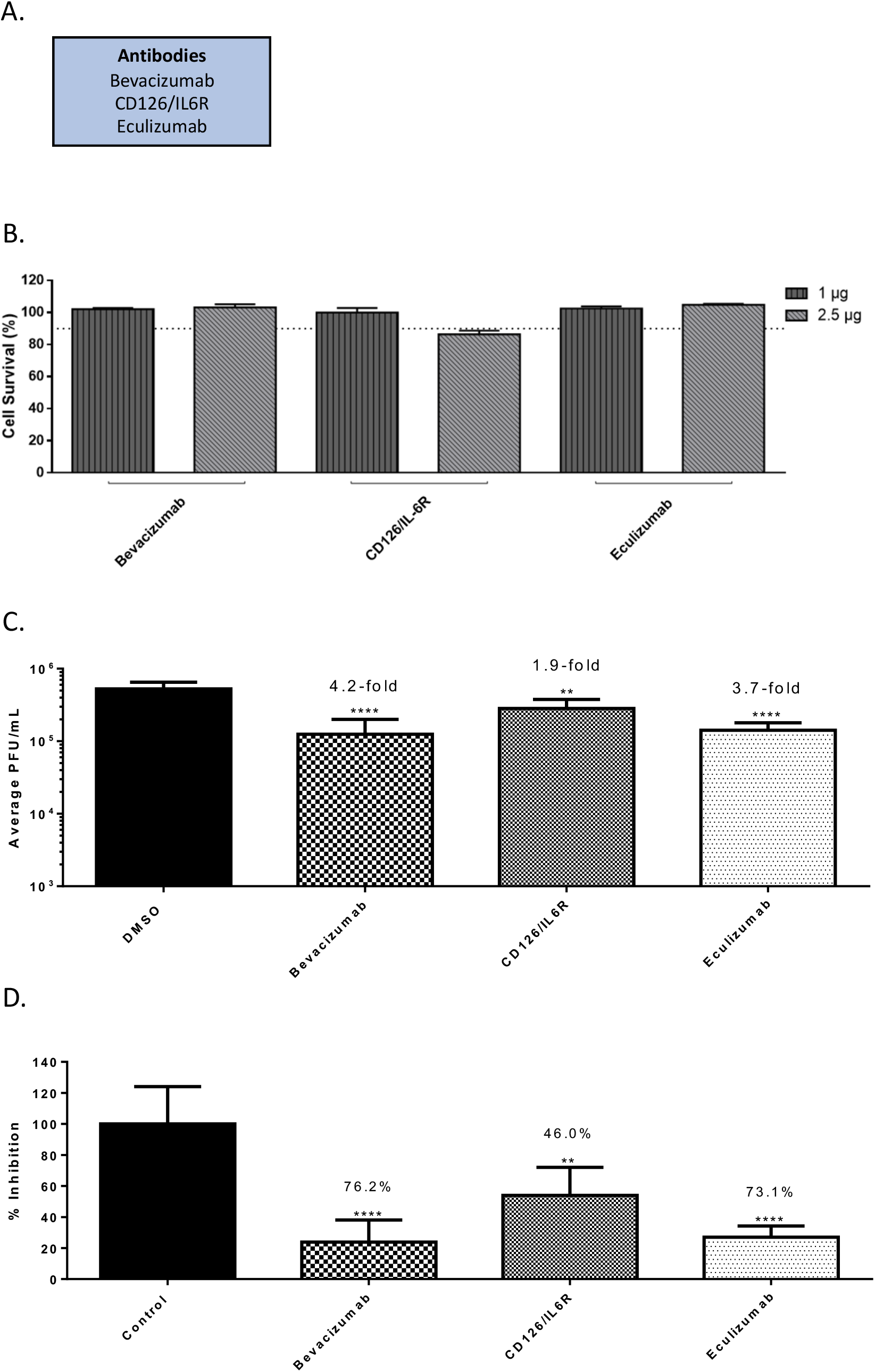
Efficacy of antibodies against SARS-CoV-2 in Vero cells 16 hpi. (A.) List of antibodies that were screened. (B.) Antibody toxicity in Vero cells at 24 hours. Cells were seeded at a density of 3E4 and overlaid with media conditioned with 0.1% DMSO or the indicated concentration of antibody. Cells were incubated for 24 hours at 37ºC at 5% CO2. Viability was determined using CellTiter Glo assay. (C.) Efficacy of antibodies in Vero cells 16 hpi. Cells were seeded at a density of 3E4 and overlaid with conditioned media containing 0.1% DMSO or maximum permissible dose of antibody. Cells were incubated for 2 hours. Conditioned media was removed. Viral inoculum (MOI: 0.1) was added to cells and incubated for 1 hour at 37ºC and 5% CO2. Viral inoculum was removed, cells were washed with DPBS, and conditioned media was returned to the cells. Culture was incubated for 16 hpi. Supernatants were collected and a viral plaque assay was used to quantify the viral titers. (D.) Inhibition that was observed at 16 hpi, represented as relative percentages. The quantitative data are depicted as the mean of three biologically independent experiments ± SD. * *p* < 0.05; ** *p* < 0.01; *** *p* < 0.001; \*\*\*\**p* < 0.0001.

### Interference of anti-SARS-CoV-2 small molecules with ACE2:Spike protein interaction

A VSV-G pseudovirus with a firefly luciferase reporter which retains the SARS-CoV-2 spike protein on the surface was used to query if the prioritized compounds affected early steps of the infectious process. The pseudovirus is capable of ACE2-based viral entry which can be quantified by measuring intracellular luciferase expression. As the pseudovirus is not capable of viral RNA synthesis, the inhibitory effect is exclusive to the early entry and post-entry steps. The stock pseudovirus was diluted 1:5 in media as specified in the protocol and was used to infect Vero cells in the presence of the inhibitor and luciferase expression was measured at 16 hours post-infection (**Figure S2A**). While Nitazoxanide exerted a 74.1% inhibition of luciferase expression, Maraviroc and FTY720 produced 28.6% and a 41.1% inhibition, respectively, of reporter expression suggesting that Maraviroc or FTY720 were likely to inhibit additional steps besides entry to achieve viral inhibition.

To further ascertain if this observed inhibition to viral entry exerted by Nitazoxanide involved interference in the molecular interaction between the viral spike protein and the cellular ACE2 receptor, a biotinylated receptor binding assay was employed. In this assay, the potential for the inhibitor to interfere with the binding of spike protein and ACE2 receptor is quantified as a measure of biotin expression. The assay involves using a nickel-coated plate to which his-tagged ACE2 is added. The drug is then added to the plate and incubated for 1 hour at room temperature after which the SARS-CoV-2 spike protein bound to biotin is added and the plate is incubated for 1 hour after which Streptavidin-HRP is added. Chemiluminescence is measured following the addition of HRP substrate which is a reflection of the interaction between the ACE2 protein and S-protein. Quantification of luminescence in drug-treated wells as compared to the DMSO-treated control demonstrated that Maraviroc, FTY 720 or Nitazoxanide did not interfere with the interaction at lower concentration of the inhibitor (**Figure S2B**). Atorvastatin elicited a modest interference in the interaction as demonstrated by a reduction of 31.7% (p<0.01) of biotin expression (**Figure S2B**). When the experiment was conducted at higher concentrations of the drug (100 μM), FTY720 was able to exert a modest interference (27% relative to control, p<0.01), while Maraviroc and Nitazoxanide treatment did not produce any impact on the interaction. The data suggest that the mechanism of inhibition for Maraviroc and Nitazoxanide is unlikely to involve disruption of receptor binding by the viral spike protein and is likely that early post-entry steps are impacted by the drug treatment (**Figure S2B)**.

### Impact of Maraviroc on intracellular events in SARS-CoV-2 infected cells

The observations that Maraviroc exhibited strong inhibition of SARS-CoV-2 when infection was performed in the presence of the inhibitor, without necessarily impacting the interaction of the viral surface spike protein with the ACE2 receptor suggested that alternate mechanisms that may be critical for the establishment of a productive viral infection may be impacted by Maraviroc. Evidence in the literature indicates that the coronavirus spike protein drives membrane fusion resulting in the formation of multinucleate cells (Fehr and Perlman, 2015; Lontok et al., 2004; Sisk et al., 2018). Notably, this phenomenon was observed to be elicited by over expressed S-protein (spike protein) as well, suggesting that it is independent of functionality from other proteins. As a next step, the impact of Maraviroc on S-protein based cellular fusion was assessed by confocal microscopy (**Figure 3A**). Vero cells were transfected with an over expression construct that expressed the SARS-CoV-2 spike protein. Transfected cells were maintained in the presence of Maraviroc for up to 48 hours to permit expression of S-protein. DMSO-treated cells were maintained as controls to ascertain baseline numbers of multinucleate cells. The cells were fixed and stained at 48 hours post transfection and the extent of formation of multinucleate cells were quantified by confocal microscopy (**Figure 3A-B**). The data demonstrate that increased expression of S-protein in Vero cells results in the formation of large multinucleate cells that typically contain 4-8 nuclei per cell. Vero cells transfected with a GFP control plasmid did not form multinucleate cells (data not shown). Interestingly, maintaining the transfected cells in the presence of Maraviroc reduced the frequency of multinucleate cell formation by approximately 2-fold as compared to the DMSO-treated controls (**Figure 3C**), as well as decreasing the average number of nuclei in these clusters (**Figure 3D**) and % fusion **(Figure 3E)** for S-protein from wild-type and D614G strains. This outcome suggests the possibility that Maraviroc treatment may result in a decrease in viral load in the cell population by preventing membrane fusion events that impact intracellular viral multiplication and spread. The mechanism of action for S-protein to prevent cell-cell fusion is not yet identified. Staining of S-protein in Maraviroc-treated cells showed increased cytosolic staining (Figure 3B), which may indicate that Maraviroc inhibits trafficking of S-protein to the extracellular surface of the cell.

**Figure 3.**
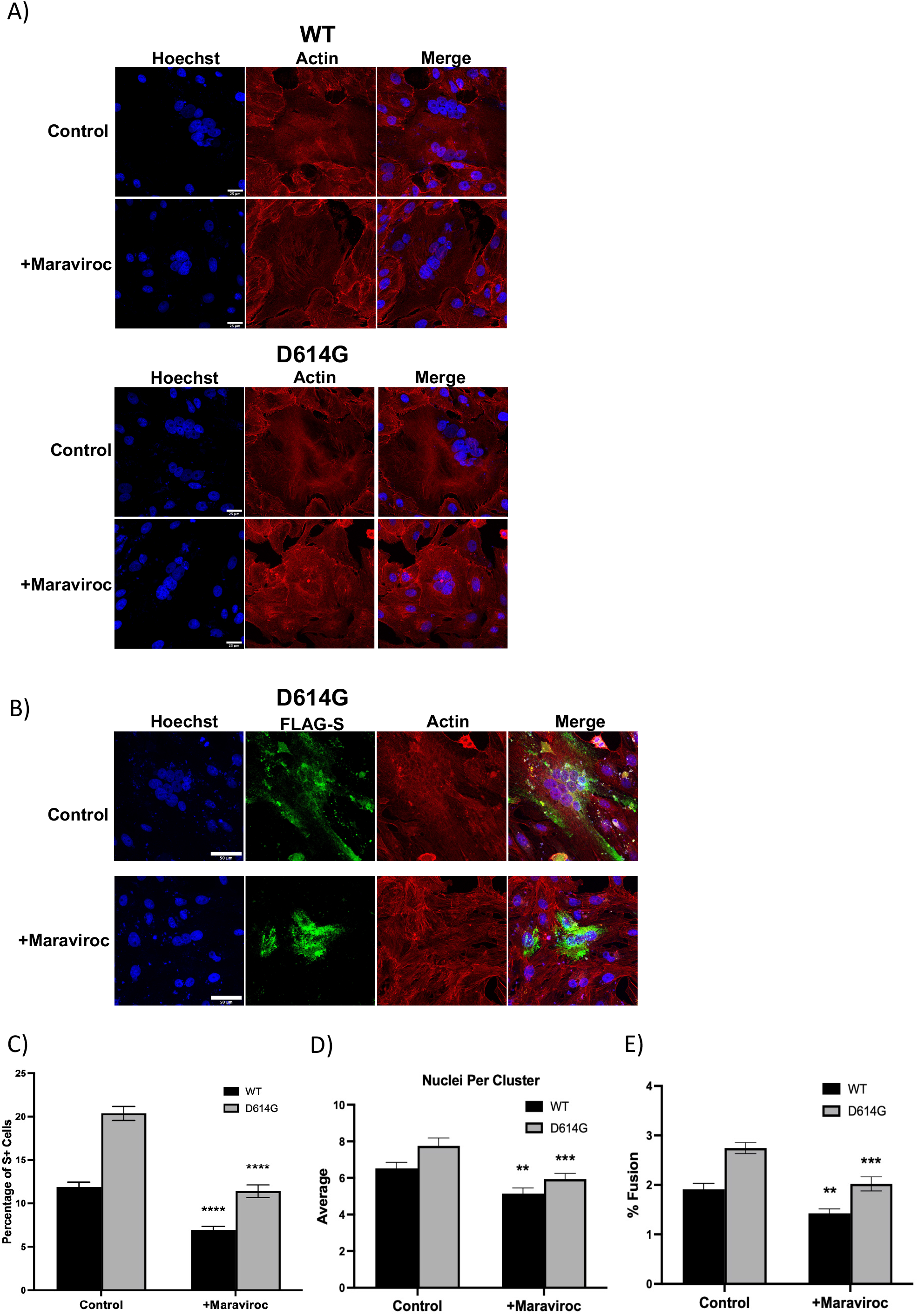
Maraviroc inhibits Wild Type (WT) and D614G strain S-protein induced cell-cell fusion. (A.) Vero cells were transfected with S protein tagged with Strep-tag II and treated with DMSO (labeled as condition ND for no drug) or 5 μM Maraviroc for 48 hours (treated immediately after transfection; labeled as condition A), or 5 μM Maraviroc for 72 hours (treated 24 hours before transfection, labeled as condition B). Immunohistochemistry showed that S protein expression resulted in the formation of multinucleated cells, which was inhibited with Maraviroc treatment. (B.) The frequency of multinucleated cells, normalized to total cell number, was reduced with Maraviroc treatment. (C.) The average number of nuclei in multinucleated cells was also reduced with Maraviroc treatment. The quantitative data are depicted as the mean of three biologically independent experiments ± Standard Error ** *p* < 0.01; *** *p* < 0.001, ***p < 0.0001.

### Determination of synergy between the anti-SARS-CoV-2 small molecules

The ability of Maraviroc to exert synergistic inhibitory outcomes when combined with FTY720, Atorvastatin or Nitazoxanide was assessed. As a first step, toxicity of the combination of drugs was assessed when the drugs were combined at 20%, 50% and 100% of their effective concentrations as observed in **Figure 1D**. To that end, the four compounds were assessed when they were combined at 1 μM each, 2.5 μM each and 5 μM each. The combinations were used to treat Vero cells and cell viability was quantified at 24 hours post treatment by Cell Titer Glo assay (**Figure 4A**). The combinations that resulted in >90% cell survival (<10% toxicity) were considered for efficacy assessment. To assess potential synergy between these compounds, infection with SARS-CoV-2 were carried out in the presence of the drug combinations and the infected cells were maintained for 16 hours in the presence of the combinations. The viral load in the supernatants were quantified at 16 hours post infection by plaque assay and relative inhibition compared to that of the DMSO control (**Figure 4B**). A modest increase in inhibitory potential was noted when Maraviroc was combined with any of the three compounds (**Figure 4C**). Inhibitory potential was increased 2.6% for FTY720, 10.6% for Atorvastatin and 11.9% for Nitazoxanide. Remdesivir was also screened for antiviral activity and possible synergy with Maraviroc. Toxicity of Remdesivir alone was below 10% for the concentrations that were screened. Combining Remdesivir with Maraviroc did not substantially increase cytotoxicity, and cell viability remained above 90%. Remdesivir alone decreased viral titer by 91.1% (p<0.0001). When Maraviroc was added to the treatment viral titers were decreased by 94% (p<0.0001).

**Figure 4.**
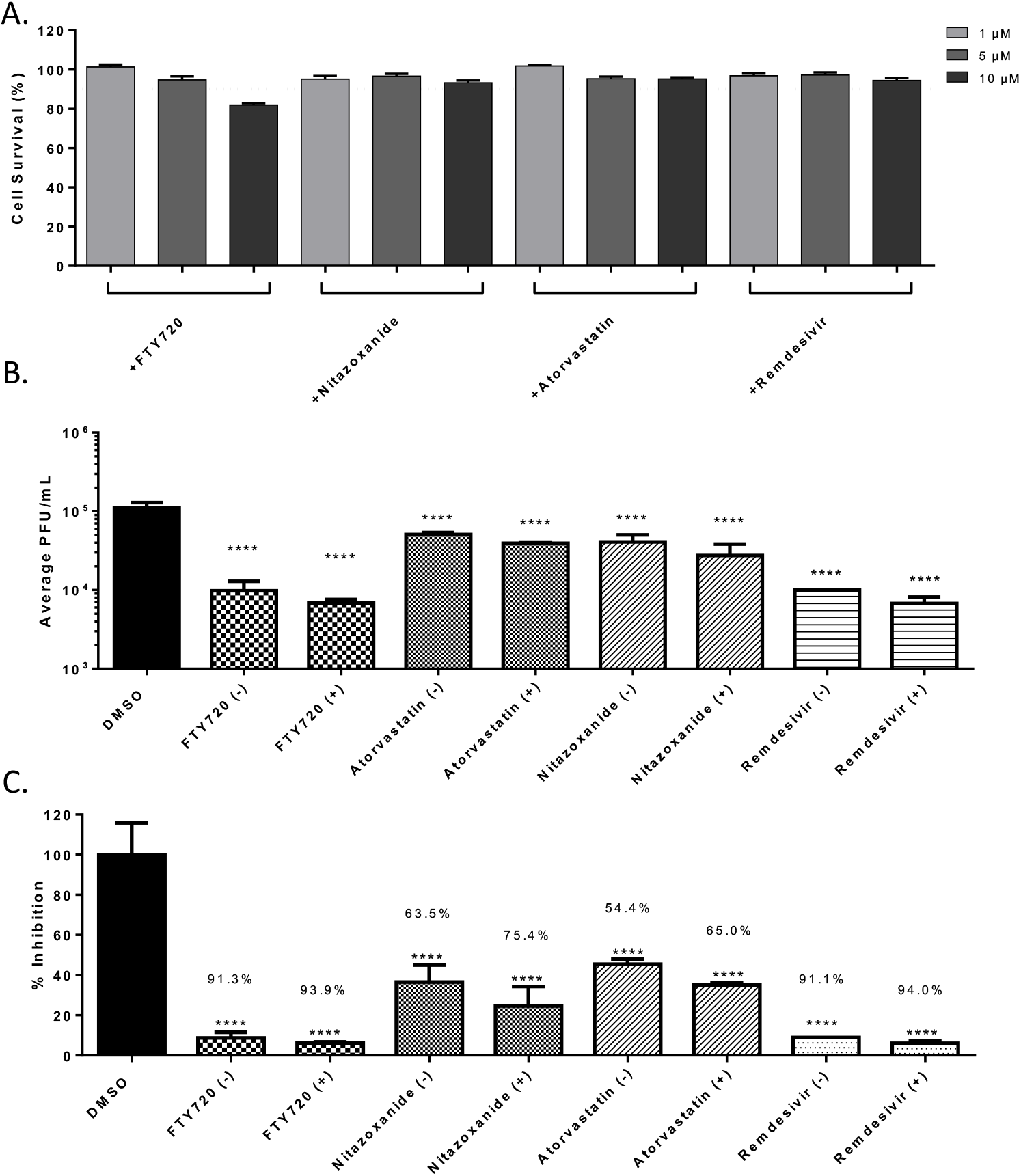
Synergistic effects of drug combinations against SARS-CoV-2 in Vero cells at 16 hpi. (A.) Combination drug toxicity in Vero cells at 24 hours. Cells were seeded at a density of 3E4 and overlaid with media conditioned with 0.1% DMSO or the indicated concentration of inhibitor. Cells were incubated for 24 hours at 37ºC at 5% CO2. Viability was determined using CellTiter Glo assay. (B.) Efficacy of inhibitors combinations in Vero cells at 16 hpi. Cells were seeded at a density of 3E4 and overlaid with conditioned media containing 0.1% DMSO or 5:5 μM of Maraviroc plus indicated inhibitor. Cells were incubated for 2 hours and conditioned media was removed. Viral inoculum (MOI: 0.1) containing 5:5 μM of inhibitor combination was added to cells and incubated for 1 hour at 37ºC and 5% CO2. Viral inoculum was removed, cells were washed with DPBS, and conditioned media was returned to the cells. Culture was incubated for 16 hpi after which supernatants were collected and a viral plaque assay was used to quantify the viral titers. (C.) Inhibition that was observed at 16 hpi, represented as relative percentages. The quantitative data are depicted as the mean of three biologically independent experiments ± SD. * *p* < 0.05; ** *p* < 0.01; *** *p* < 0.001; \*\*\*\**p* < 0.0001.

### Modeling of Maraviroc mediated inhibition of SARS-CoV-2 in a multicellular environment

To study the dynamics of SARS-CoV-2 spread under control and Maraviroc-inhibited conditions, we adapted a computational model of SARS-CoV-2 (Getz et al., 2020) to include cellular fusion driven by surface-bound spike protein, and integrated a mixed-effect pharmacodynamic response model that can simulate any combination of inhibiting cell fusion, viral entry and exit, and replication. After verifying that single-effect pharmacodynamics models that solely inhibit cell fusion or viral entry/exit cannot simultaneously match our experimental measurements and observations that Maraviroc reduces both viral load at 16 hours, and cell fusion at 48 hours, we fitted a mixed-effect model that inhibits both cellular fusion and viral entry and exit. (See **Fig. S3** and Simulation **model fitting** above.) In particular, we tuned the viral binding, endocytosis, replication, exocytosis, and pharmacodynamics parameters to match the viral titer at 16 hours, and the number of nuclei per cell at 48 hours under no treatment and at 5 μM of Maraviroc.

The fitted model suggests that exocytosis is not fully inhibited at 5 μM: because the original 3000 virions are quickly bound and endocytosed, replication and exocytosis are required to explain the presence of unbound virus at 16 hours. Moreover, because Maraviroc does not inhibit SARS-CoV-2 binding to ACE2 receptors on uninfected cells, sustained replication and exocytosis are required to outpace binding to ACE2 receptors. In spite of incomplete inhibition, fewer cells are infected by 16 hours, and fewer infected cells have a high viral load of assembled virions. (**Figure S3)**

When we extended the simulations to 2 days (**Figure 5A**), we found that the 5 μM dose **(Figure 5B**) delayed but did not prevent the spread of the infection. By 2 days, all viable cells were infected with high viral load (non-blue cells), but cell survival was increased over the untreated control **(Figure 5A)**. Increasing the simulated dose to 7.5 μM **(Figure 5C)** or 10 μM **(Figiure 5D)** further increased cell survival, and some cells remained uninfected at 2 days.

**Figure 5:**
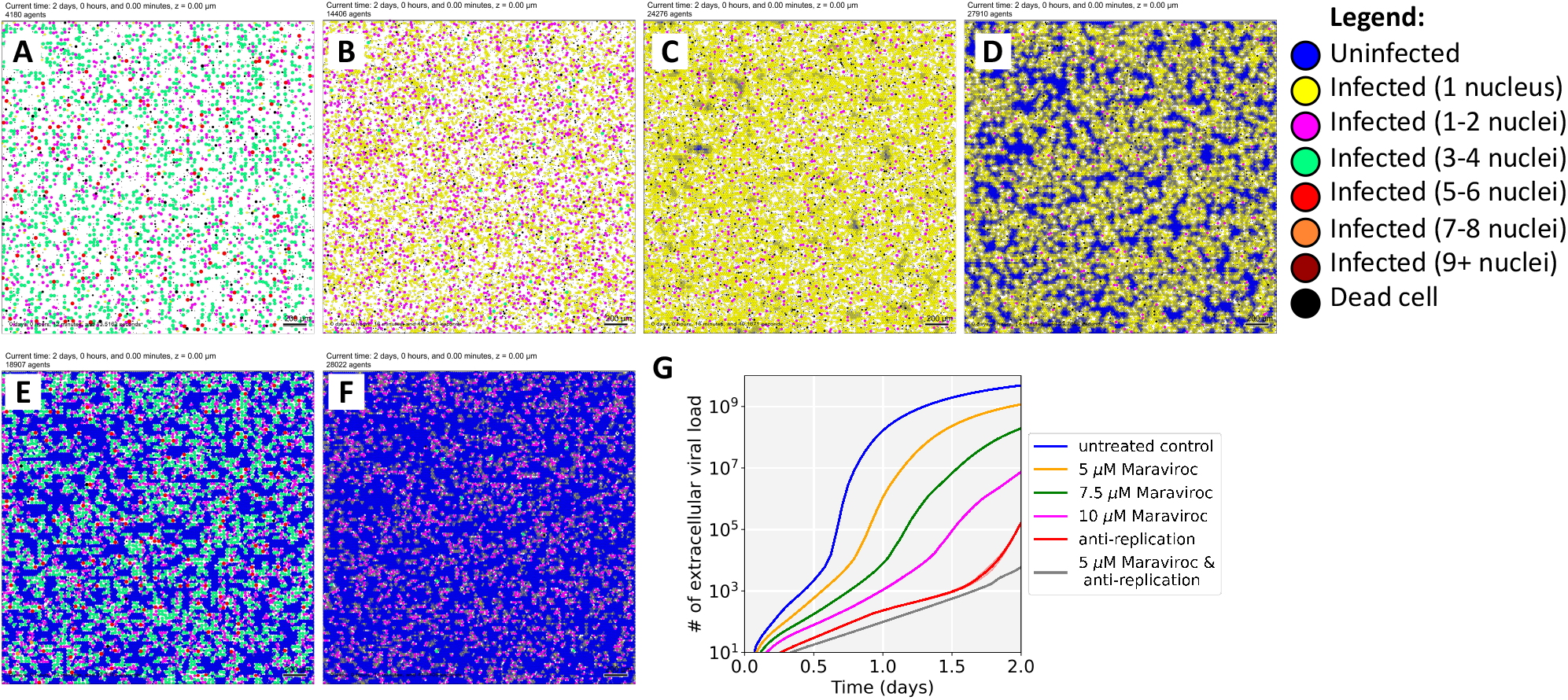
Simulation results. (A-D): We extended our *in vitro* experiments to 2 days for untreated control (A), 5 μM Maraviroc (B), 7.5 μM Maraviroc (C), and 10 μM Maraviroc (D). Note that dose escalation increases the number of uninfected cells reduces cell fusion, the number of nuclei in fused cells, and overall viral load. (E-F): We simulated a 2-day treatment with an anti-replication drug (e.g., Remdesivir) calibrated to approximately match experimental responses at 2 days (E). Without additional fitting, we then simulated a combination treatment with 5 μM Maraviroc and the same anti-replication treatment (F). Just as in the experiments, the computational model predicts an additional synergistic improvement in cell survival. We plot the impact on viral load for these virtual experiments in (G)

We also examined the potential to combine a moderate (5 μM) dose of Maraviroc with an anti-replication drug such as Remdesivir. First, we calibrated an anti-replication pharmacodynamic response to match the Remdesivir-only experiments, and simulated to 2 days (**Figure 5E**). Next, we simulated a combination of Remdesivir with 5 μM Maraviroc without any additional calibration. Just as in experiments, the simulation model predicted that the combination would provide a modest improvement (**Figure 5F**).

After verifying that the computational model recapitulated these results, we further investigated the role of cellular fusion in viral spread in untreated conditions. When we completely disabled cellular fusion in untreated conditions, we found virtually no reduction in viral load (**Figure S4A**), suggesting that extracellular spread of virions may ordinarily be more efficient at transmission than fusion. When we reduced extracellular virion spread (e.g., as might be expected in a tissue where virions are quickly removed by flow or neutralizing antibodies), we found that fusion substantially increased viral transmission (**Figure S4B-C**). Thus, cellular fusion could play a role as a “backup transmission” method when extracellular transmission is impaired by treatment or an adaptive immune response.

## Discussion

SARS-CoV-2, the causative agent of the ongoing COVID-19 pandemic, continues to have a devastating effect globally, increasing the morbidity and mortality burden on a daily basis and giving rise to significant long-term health impacts even in cases of mild infections (Hayes et al., 2021; Michelen et al., 2021; Nalbandian et al., 2021). From the perspective of bringing quick treatment options to the clinic, there continues to be heightened focus on repurposing FDA-approved drugs. When a therapeutic intervention approach to treat COVID-19 is to be evaluated, it cannot be ignored that the population that presents itself at the clinic is likely to fall in the more severe disease scale, with a high probability of co-morbidities and ongoing medication regimens. This possibility has to be considered from two angles: 1. the aggressive therapeutic strategy that may be used to treat SARS-CoV-2 infection should not interfere with ongoing drug regimens or exert unforeseen catastrophic outcomes; 2. the possibility of treatment strategies that are commonly used to treat disease conditions such as cardiovascular disease, asthma or immune system disorders may be used to also treat SARS-CoV-2 infection. The latter aspect offers not only the possibility of being able to safely use some of these therapeutics in the treatment of these high risk groups, but also paves the way for determining if any of these ongoing therapeutic regimens may offer a level of protection to the target group by preventing escalation of disease severity.

In this study, a total of 22 small molecules and biologics that are commonly used to treat disease states including viral, bacterial and parasitic infections, heart disease and inflammatory conditions such as multiple sclerosis were analyzed for potential antiviral activity against SARS-CoV-2 in a cell culture model. The investigation identified four small molecules that demonstrated statistically significant anti-SARS-CoV-2 activity, namely, Maraviroc, FTY720, Nitazoxanide and Atorvastatin. Maraviroc elicited a statistically significant and consistent inhibitory effect against SARS-CoV-2 **(Figure 1C**). Maraviroc is an inhibitor of the cellular CCR5 receptor and is used to treat HIV infection. Interestingly, Maraviroc is under consideration for a clinical trial as a COVID-19 therapeutic as it is known to be well tolerated in human populations, even in the context of a compromised immune system (ClinicalTrials.gov Identifier: NCT04435522). The rationale behind the use of Maraviroc is that it would be beneficial in preventing immune cell depletion (or exhaustion) and increasing routing of inflammatory cells away from sites that sustain inflammatory damage during COVID-19 disease, thus additively contributing to lowering viral load and inflammatory damage in organs such as the lungs.

FTY720 (Fingolimod) is a drug that is used to treat multiple sclerosis which demonstrated robust inhibitory activity against SARS-CoV-2 (**Figure 1C**). It functions as an immune modulator that is currently undergoing clinical trials for use as a COVID-19 therapeutic (ClinicalTrials.gov Identifier: NCT04280588). The rationale for consideration of FTY720 is that it may be able to reduce the inflammatory manifestations of ARDS in the lung by decreasing pulmonary edema and hyaline membrane formation, possibly if used in conjunction with additional methods including ventilator support. Furthermore, the mechanisms of FTY720-mediated reduction of egress of autoreactive lymphocytes from lymphoid tissues may play additional roles in decreasing systemic inflammatory damage. FTY720 treatment may produce short term fluctuations in cardiovascular parameters and blood pressure due to agonistic and antagonistic impacts on Sphingosine-1-phosphate receptor (S1PR) which is differentially expressed in tissues including cardiac cells (Camm et al., 2014). FTY720 has also been implicated in cases of Fingolimod associated macular edema, especially when administered in the same regimen as for multiple sclersis (Pul et al., 2016). These factors will require consideration if FTY720 is to be evaluated for COVID-19 treatment.

Nitazoxanide, an FDA-approved drug used to treat diarrhea, exerted antiviral activity in the context of SARS-CoV-2 infection (**Figure 1C**). There have been recent published reports that refer to the use of Nitazoxanide in combination with other drugs including Azithromycin and Ivermectin, both of which were demonstrated to show anti-SARS-CoV-2 activity in cell culture (Kelleni, 2020). Nitazoxanide has been demonstrated to have antiviral activity against MERS-CoV and potentiate interferon responses (Rossignol, 2016). In the context of MERS-CoV infection model, Nitazoxanide was also shown to decrease the expression of the proinflammatory cytokine IL6, which is also implicated in the inflammatory damage elicited due to SARS-CoV-2 infection. Nitazoxanide is under consideration for a clinical trial as a COVID-19 treatment strategy (ClinicalTrials.gov Identifier: NCT04435314).

Atorvastatin, a commonly used small molecule for the treatment of cardiovascular disease demonstrated modest antiviral activity against SARS-CoV-2 when compared to the other effective compounds (**Figure 1C**). Recently published outcomes suggested that statin use in patients is likely to be linked to decreased mortality, ventilator use and hospitalizations in the case of COVID-19 patients (Zhang et al., 2020). Atorvastatin is currently undergoing clinical trials as adjunctive therapy for potentially addressing the cardiovascular complications associated with COVID-19 (ClinicalTrials.gov Identifier: NCT04380402).

Studies performed by varying the infection condition, specifically the presence or absence of the candidate drug at the time of infection revealed that including the drug during infection produced a higher rate of inhibition (data not shown). This observation led to the hypothesis that the candidate compounds, Maraviroc, FTY720, Nitazoxanide and Atorvastatin may interfere with entry and/or early post-entry steps in the SARS-CoV-2 infectious process. To investigate this possibility further, a VSV-G pseudovirus which expresses the SARS-CoV-2 spike protein on the surface was used in the context of these inhibitors. The extent of inhibition observed with the pseudovirus was significantly lower than what was noted in the context of SARS-CoV-2 (**Figure S2A**) suggesting that additional steps beyond viral entry are likely to be influenced by these small molecules. It may also be possible that the VSV glycoproteins that are also expressed on the surface of the pseudovirion may interfere with the inhibition that these compounds may exert on the SARS-CoV-2 spike protein. Nevertheless, the observation that the early entry steps are impacted by these compounds led to the question whether they interfered with the interaction between the viral spike protein and the host ACE2 receptor. Analysis of binding interaction by a reporter assay in the presence and absence of the inhibitor revealed that while Atorvastatin may elicit a modest level of interference in the interaction between the spike protein and ACE2, Maraviroc and FTY720 did not, even at higher concentrations (**Figure S2B**). Maraviroc and FTY720 were the most potent of the inhibitors that were tested in this effort and the data supported the idea that the mechanism of inhibition is not likely to involve interference with receptor binding.

Coronavirus spike protein has been shown to be involved in membrane fusion, both in the context of fusion between viral and host cell membranes and in the context of host cell membranes (Tang et al., 2020). The membrane fusion event is believed to be mediated by the S2 domain of the extracellular stalk portion of the spike protein. Within the S2 domain, a short, loosely defined fusion peptide, a stretch of 15-25 amino acids is important to initiate the fusion event and is considered to be very sensitive to mutations (Madu et al., 2009). The SARS-CoV-2 fusion peptide region in S2 is considered to be 93% similar to that observed in SARS-CoV-1, thus indicating a high degree of conservation in the fusion peptide (Tang et al., 2020). SARS-CoV-1 S-protein was shown to initiate cell fusion between human cells, which was also observed in the case of SARS-CoV-2 (Ou et al., 2020) and the protease TRPRSS11a is thought to aid this fusion event because of S-protein processing in the context of SARS-CoV-1 (Kam et al., 2009). Studies that were conducted with an over expressed SARS-CoV-2 S-protein expression construct demonstrated an increased incidence of multinucleate cells with an average of 4-7 nucleic per multinucleate cell (**Figure 3)**. When the transfected cells were maintained in the presence of Maraviroc, a statistically significant reduction in the number of multinucleate cells were observed (**Figure 3B**). This observation raises the interesting possibility of viral spread in a multicellular environment mediated by cell fusion. Such a possibility holds significant relevance to the efficacy of convalescent serum use as an intervention strategy because the intracellular viral population will be shielded from the antibodies and will continue to persist. Synthetic antibodies or nanobodies that are membrane permeable may prove to be effective in such cases. If such a possibility is true, it will explain the observed cases of “relapse” or “prolonged persistence” of viral RNA even in individuals who do not exhibit any disease symptoms. Such an event will also have a significant impact on vaccine design, particularly those that rely on humoral responses to extracellular virus, drawing attention to CD8 mediated responses as being an important requirement for an efficacious vaccine.

An interdisciplinary coalition is actively developing an open source, multiscale simulation model for SARS-CoV-2 infection dynamics in tissue (Getz et al., 2020), which can model the heterogeneous dynamics of individual cells and the spread of viral particles through a tissue (or cell culture). This study adapted and extended the simulator by adding celllar fusion and a Maraviroc pharmacodynamics model to each cell and calibrating to the untreated control and 5 μM experiments at 16 hours. As employed here, the simulator emphasizes the early entry and post entry events in the context of Maraviroc treatment as a test case with a goal of implementing a full *in vivo* simulation model to assess the clinical applicability of the drug. The simulations revealed that the inhibition is likely to extend protective effects in a multicellular setting and decrease cell death in relation to a lowered viral load if the effective drug concentration can be maintained. Because the model explicitly simulates viral binding, endocytosis, replication, release and spread as well as cellular fusion, it can help “fill in the gaps” between limited experimental observations. In this effort, the model suggests that antiviral drugs such as Maraviroc can reduce viral titer by over two orders of magnitude even with partial inhibition of its target at moderate doses. Moreover, it also agrees with experimental findings that combining Maraviroc could complement existing drugs that target replication (e.g., Remdesivir) for further efficacy, and that targeting cellular fusion may be beneficial when contact-based viral transmission otherwise evades treatment. Future simulation studies will assess whether Maraviroc can act synergistically with innate immune responses to improve control of the virus and reduce tissue damage. While simulations were employed in the context of Maraviroc as an initial demonstration of utility, the additional inhibitors identified in this study will be queried independently for control of viral and inflammatory load, relative pharmacodynamics properties as related to antiviral and anti-inflammatory activities and synergistic outcomes. The simulation model could also afford the opportunity to assess the relative impact of cell-to-cell viral transmission (e.g., via fusion) compared to diffusive dissemination of endocytosed virus particles, and whether treatments like Maraviroc could ultimately halt these alternative forms of viral spread in tissues.

In conclusion, our data demonstrate that Maraviroc, FTY720, Nitazoxanide and Atorvastatin are viable candidates that may be used to treat individuals infected with SARS-CoV-2. These drugs are commonly used to treat a wide demographic of the human population and are known to be well tolerated in humans. The data presented in this effort will also provide support to the ongoing clinical trials that are exploring the utility of these compounds as COVID-19 treatment strategies.

## Acknowledgements

The following reagent was deposited by the Centers for Disease Control and Prevention and obtained through BEI Resources, NIAID, NIH: SARS-Related Coronavirus 2, Isolate USA-WA1/2020, NR-52281. We would like to thank Dr. Frederick Holtsberg for kindly providing us the VSV-G pseudovirus. This research was supported by GMU startup funds to AN; NIH R35GM119617, NSF CMMI-1653299, and VCU COVID-19 Rapid Research Funding program to DC, and by generous support from the Jayne Koskinas Ted Giovanis Foundation for Health and Policy (PM and YW). YW and PM acknowledge access to FutureSystems at the Digital Science Center, Luddy School of Informatics, Computing, and Engineering, and to the Big Red 3 supercomputer at Indiana University, Bloomington. We would like to thank all members of the Narayanan laboratory for help with manuscript review.

## Author Contributions

Conceptualization, K.R., K.T., Y.W., M.G., S.N., D.C., P.M. and A.N.; Methodology, K.R., K.T., Y.W., M.G., D.C., P.M. and A.N.; Software, Y.W., M.G. and P.M.; Formal Analysis, K.R., K.T. and Y.W., M.G.; Investigation, K.R., K.T., Y.W., M.G., A.B., F.A. and N.B.; Resources, K.R., K.T., Y.W., M.G., A.B., F.A., N.B., S.N., D.C., P.M. and A.N; Writing – Original Draft, K.R., K.T., Y.W., M.G., S.N., D.C., P.M. and A.N.; Writing – Review & Editing, K.R., K.T., Y.W., M.G., A.B., F.A., N.B., S.N., D.C., P.M. and A.N.; Visualization, K.R., K.T., and Y.W.; Supervision, A.N., D.C. and P.M.; Funding Acquisition, A.N., D.C., P.M. and S.N.

## Declaration of Conflict of Interests

The authors declare no conflict of interest with any aspect of this work.

## Notes

### Competing Interest Statement

The authors have declared no competing interest.

### Summary of Updates

This include improved analyses of experiments, better contextualization, refined simulation results, and updated discussion.

